# Historical abundance and distributions of *Salpa thompsoni* hot spots in the Southern Ocean, with projections for further ocean warming

**DOI:** 10.1101/496257

**Authors:** Angelika Wanda Słomska, Anna Panasiuk, Agata Weydmann-Zwolicka, Justyna Wawrzynek-Borejko, Marta Konik, Volker Siegel

**Affiliations:** University of Gdansk, Department of Marine Plankton Research, Faculty of Oceanography and Geography, 46 M. J. Pilsudskiego Avenue, Gdynia, Poland; University of Gdansk, Department of Marine Ecosystem Functioning, Faculty of Oceanography and Geography, 46 M. J. Pilsudskiego Avenue, Gdynia, Poland; The Institute of Oceanology of the Polish Academy of Sciences, 55 Powstancow Warszawy Street, Sopot, Poland; Sea Fisheries Institute, 9 Palmaille, 22767 Hamburg, Germany

**Keywords:** climate change, environmental impact assessment, invertebrates, invertebrates, ocean, zooplantkon

## Abstract

1. Over the last three decades, a significant variability in *Salpa thompsoni* occurrence has been observed as a response to the environmental fluctuations of the Southern Ocean ecosystem, e.g. changes in sea surface temperature as well as shrinking of ice-cover extent around the cold Antarctic waters.
2. This study presents the historical data of salps abundance from the southwest Atlantic Sector of the Southern Ocean and covers time span of 20 years. Presented dataset allowed to track previous fluctuations in Antarctic salp abundance and enabled to combine their distribution with different bottom depth, thermal and ice conditions. The subsequent goal of this work was to reveal hot spots of salps location and to predict the future range of *S. thompsoni* distribution with upcoming climate warming in the next 50 years.
3. Results of our study revealed that the highest salp number was located mostly in the shallow shelf waters with ice-cover and lower temperature. In the studied area, *Salpa thompsoni* hot spot distributions have been located mostly around Elephant Island but also within islands around Brensfield and Gerlache Straits, as well as to the south near the cold Bellingshausen Sea. The inference of future salp distribution demonstrated that the range of *S. thompsoni* would presumably move southwards enlarging their habitat area by nearly 500 000 km^2^.

## Introduction

Over the last 30 years, the rise of atmospheric greenhouse gas has influenced the average global atmospheric temperature, with an increase of 0.2°C, and the mean global sea surface temperature (SST) increasing by approximately 0.4°C (Levitus, Antonov & Boyer, 2005; Levitus, Antonov, Boyer & Stephens, 2000). Ocean heating is particularly pronounced in the most vulnerable areas which include the Southern Ocean, especially in the Antarctic Peninsula region (Whitehouse, Meredith, Rothery, Atkinson, Ward & Korb, 2008; Zwally, Comiso, Parkinson, Cavalieri & Gloersen, 2000). Despite this, study by Turner et al. (2016) based on the stacked air temperature record since the 1990s, has suggested that there was no evidence of climate warming in the mentioned area. However, hydrological processes and modifications recorded in the Southern Ocean are much more complex. It should be recalled that clear proof of decreasing sea ice extent in the Bellingshausen and Amundsen Seas has been numerously recorded, and this phenomenon could be tied to continuous ocean heating (Cavalieri & Parkinson, 2012). In particular, summer water temperature along the Western Antarctic Peninsula increased by 1.3°C over 50 years (Meredith & King, 2005), and near South Georgia a 1°C increase has been recorded over the last 80 years (Whitehouse, Meredith, Rothery, Atkinson, Ward & Korb, 2008). The Weddell Deep Water (WDW) has warmed by ∼0.032°C per decade, which is similar to the temperature changes around South Georgia and in the entire Antarctic Circumpolar Current (0.03–0.07°C per decade) (Nicol, Croxall, Trathan, Gales & Murphy, 2007; Whitehouse, Meredith, Rothery, Atkinson, Ward & Korb, 2008).

The current phenomenon of environmental stress caused by climatic and anthropogenic pressures leads to shifts of frontal hydrological zones, existing habitats, and is also likely to modify structure, spatial range and seasonal abundance of Antarctic key species (Atkinson, Siegel, Pakhomov & Rothery, 2004; Loeb & Santora, 2012; Pakhomov, Froneman & Perissinotto, 2002; Richardson, 2008;; Ross et al., 2014; Steinberg et al., 2015). The geographical range of crucial Antarctic taxa can move southwards to remain in optimal thermal conditions, squeezing their distribution range closer to the Antarctic continent (Atkinson, Siegel, Pakhomov & Rothery, 2004; Loeb & Santora, 2012; Richardson, 2008; Ross et al., 2014). At the same time, more favorable habitats can be opened for more flexible organisms that are capable of adaptation like gelatinous salps (Goodall-Copestake, 2016; Jue et al., 2016; Mackey et al., 2011; Ross et al., 2014; Steinberg et al., 2015).

Antarctic salp *Salpa thompsoni* (Foxton, 1961), together with the krill *Euphausia* superba (Dana, 1850), are among the most important filter-feeding species of the Southern Ocean, but only Antarctic krill is considered as a major food source for many top predators, including fish, penguins, seals and baleen whales in a very short Antarctic trophic chain (Loeb & Santora, 2012). Therefore, regarding the current threat to decreasing krill numbers, and due to a changing climate conditions, there is a growing concern that salps may locally replace krill. Possible mechanisms underlying these observations include the southerly movement of the sea ice edge, water temperature fluctuations, reconstruction of the phytoplankton structure, and changes in Antarctic Circumpolar Current transport pathways. All these factors might cause shifts in Antarctic krill populations, and create a free ecological niche available for pelagic tunicates.

*Salpa thompsoni* is typically oceanic, has an extensive circumpolar distribution (45-55°S) (Foxton, 1966; Pakhomov, Froneman & Perissinotto, 2002), is not ice-dependent like *Euphausia superba* and it is usually found in areas with lower food concentrations (Atkinson, Siegel, Pakhomov & Rothery, 2004; Loeb et al., 1997; Pakhomov, Froneman & Perissinotto, 2002;Siegel & Loeb, 1995; Steinberg et al., 2015). Antarctic salps prefer water masses of higher temperature and lower productivity across the Circumpolar Current, therefore the highest salp abundance is observed along with the absence or at least really low number of Antarctic krill (Atkinson, Siegel, Pakhomov & Rothery, 2004; Siegel & Loeb, 1995). Previous studies provided some evidence that the greatest changes in abundance and distribution of salps in the Atlantic Southern Ocean are associated with the sea ice loss, transitional periods between El Niño-La Niña (ENSO), and with the shifts of the Southern Boundary (SO) of Antarctic Circumpolar Current (ACC) (Atkinson, Siegel, Pakhomov & Rothery, 2004; Chiba, Ishimaru, Hosie & Wrigh, 1999; Pakhomov, Froneman & Perissinotto, 2002; Siegel & Loeb, 1995).

The primary focus of these presented studies was to identify the main barriers hindering the salps occurence and provide missing predictions of the salp population dynamics. Based on the historical point measurements of *Salpa thompsoni* abundance in the Atlantic Sector of the Southern Ocean within over twenty years of time span (1975-2001) a spatial distribution of salp populations was reconstructed. Using the Hot Spot analyses, it revealed the environmental conditions controlling their propagation in the investigated area with an insight into the upcoming 50 years in the age of global warming.

## Methods

### Study area

The study area covered the region of the Western Antarctic Peninsula and South West (SW) Atlantic sector of the Southern Ocean (Fig. 1).

**Fig. 1.**
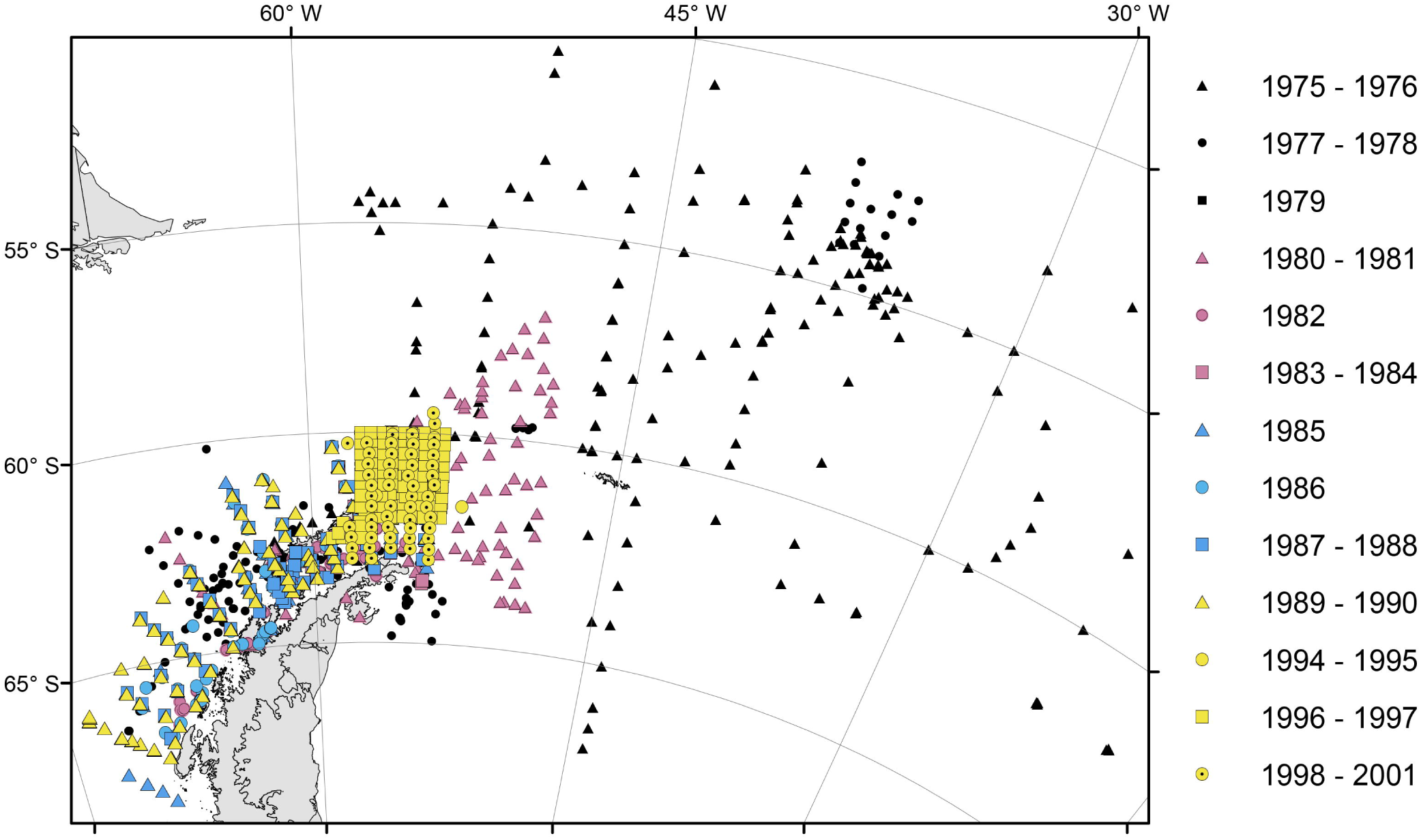
The locations of the sampling stations with the year of the cruise depicted in different color and shape.

The waters flowing around the Antarctic Peninsula are a specific and unique mixture of various water masses (Martinson et al., 2008). Here we can observe the mixing of warm, nutrient rich Circumpolar Deep Water (CDW) with cold, Antarctic shelves and coastal water masses, which have important effects on physical and biological processes (Ducklow et al., 2012). Antarctic Peninsula (AP) water mass distribution is restricted by bathymetric features such as continental slope, coastal or shelf regions (Martinson, Stammerjohn, Iannuzzi, Smith & Vernetd, 2008). This region is also more susceptible to the influence of the Antarctic Circumpolar Current (ACC) than other regions around the Antarctic where the ACC is rather distant from the shelf, typically separated by a polar gyre (Martinson, Stammerjohn, Iannuzzi, Smith & Vernetd, 2008). The marine ecosystems of the studied area are characterized by high diatom concentrations and consequently by a high primary production, krill predominance and a variety of higher vertebrate consumers (Siegel, Skibowski & Harm, 1992). Macronutrient concentration, such as nitrate and phosphate, are generally within the AP region compared to more open waters, due to three factors - high concentrations in deep water, winter deep water mixing resupplying the surface layer after biological depletion, and micronutrient (iron) limitation (Ducklow et al., 2012).

### Sampling

Samples were collected during the German and British research surveys (RV ‘Walther Herwig’, ‘John Biscoe’, ‘Polarstern’ and ‘Meteor’ cruises) using a Rectangular Midwater Trawl (RMT8) (mesh size 4.5 mm, mouth opening of 8 m^2^) (Baker, Clarke & Harris, 1973) between 1975 and 2001, during early summer/late autumn. The RMT8 was equipped with a real-time depth recorder (TDR) and sampled from the upper 200 m of the water column or 10 m above the bottom, in shallow areas. A double oblique net tow was carried out routinely at all stations with a standard station grid at a tow speed of 2.5 to 0.5 knots. Calibrated flowmeters, mounted on the net frame, were used to estimate the volume of filtered water during each haul. Filtered water volume was calculated using equations from Pommeranz, Hermann & Kühn (1992). All samples from the surveys were processed at sea. Such approach led to collecting 1872 samples in total, out of which the salps were removed immediately after the tow, counted prior to other sample processing and were stored in 4% buffered formaldehyde for later measurements. Water temperature (T) and salinity (S) were measured prior to sampling (upper 200 m), however this data was available only for a limited number of samples: T for 461 samples and S for 212 samples. Ice coverage was estimated from the bridge of the vessel by visual observation and by checking the radar, to assess the ice extended for more than one mile around the vessel. The Ice coverage was expressed in the % coverage to four classifications 0, 1, 2 and 3, where 0 means no ice, 1 <15%, 2 < 50% and 3 > 50 % of ice cover within one nautical mile.

### Reanalysis datasets

Temperature and salinity were not measured during all sampling surveys therefore we compensated our dataset for years 1970 – 2001 using available complete monthly product of the Sea Surface Temperature (SST). Due to the fact that in the Antarctic area persistent cloud cover limits available satellite information we used results of reanalyses HadISST1 published by the Met Office Hadley Centre for Climate Prediction and Research. The continuity of the SST maps had been achieved due to use of the Reduced Space Optimal Interpolation (RSOI) technique. The time series covered ranges from 1871 to date and it was based on the *in situ* observations. However, data collected in the Met Office Marine Data Bank (MDB) had been quality controlled e.g. corrected for bias and gridded onto a 1° area grid. Auxiliary data were included in order to increase quality of the product such as the monthly median SSTs from the Comprehensive Ocean-Atmosphere Data Set (COADS) which were incorporated in to the HadISST1 product for years 1871 – 1995. Since 1982 information from the satellite AVHRR sensor has also been incorporated into the dataset in order to significantly increase spatial data coverage and thus product quality (Rayner et al., 2003).

Information about the sea ice cover was obtained from the National Snow and Ice Data Center (NSIDC). The ‘Sea Ice Extent’ product (ID: G02135) included in this study was downloaded in the form of complete time-series from the November 1978 to the present. It was created at the Goddard Space Flight Center (GSFC). The information about sea ice was derived from passive microwave satellites the Nimbus-7 Scanning Multichannel Microwave Radiometer (SMMR), the Special Sensor Microwave/Imager (SSM/I) since July 1987, and the Special Sensor Microwave Imager/Sounder (SSMIS) instruments onboard satellites launched as part of the Defense Meteorological Satellite Program (DMSP). The data covered the Southern Hemisphere and were bounded on the north by the parallel of 39.23° southern latitude at spatial resolution of 25 km × 25 km (Fetter et al., 2017). The sea ice presence was expressed in a dichotomic (binary) way, where: 1 refers to sea ice cover present and 0 refers to no sea ice station. The threshold for the ice sheet was adopted as the point, where the average sea ice concentration in that month drops below 15 percent which is consistent with the operational definition presented in the IPCC reports (Vaughan et al., 2013; Fetterer et al., 2017).

### Data analysis

In order to test possible correlations between significant environmental variables, the nonparametric Spearman’s rank-order correlations were calculated prior analyses. Due to the log-normal distribution of the data, further analyses were performed on the log-transformed abundance data [x’=log (x+10)] using STATISTICA 12.0 PL (Statsoft Inc.) software. To reveal if the sea ice concentration both during sampling and in the preceding winter season (reanalysis dataset) had any impact on the number of salps during the closest Antarctic summer, the analysis of variance ANOVA (Kruskal Wallis test) and a probit regression model was used, respectively. Additionally, the Generalized Additive Models (GAMs, Poisson distribution) implemented in CANOCO 5 software were used to examine the response of salps abundance by determining their most probable spatial distribution in relation to sea surface temperature, bottom depth, and salinity, which were tested separately due to the varying number of available in situ measurements of bottom depth, T and S (1872, 461 and 212 samples, respectively).

### Spatial analyses

The Global Moran’s I statistic was used to test whether data are dispersed or there is a similarity between the collected samples. It was chosen, because The Global Moran’s I Index assesses the spatial autocorrelation based on values as well as features locations. Z-score statistics and p-value are computed along with the index in order to estimate the significance of the result. A positive Moran’s I Index value proves existence of the clusters, whereas a negative shows tendency towards dispersion (Moran, 1950). As the statistics pointed to strong clustering of the data (z-score, representing measures of standard deviation: > 60, p-value: <0.01) they were divided into three subsets based on the decadal time periods. The division was based on spatial distribution of the sampling points, as the research stations were placed over a vast area, and were confirmed as being statistically significant (z-score: >98, p-value: <0.01). Therefore, both ice cover interpolation and hot spot analysis were applied separately to each decade ‘70s, ‘80s, and ‘90s (including year 2001, because of its similarity to the cruises from 94/95 and 96/97). In order to interpolate ice coverage, the Kernel Interpolation with Barriers method was performed. The coastline of the Antarctic Peninsula from Antarctic Digital Database (ADD) was used as a barrier in the analysis. The kernel function was set to first order polynomial with the ridge parameter retaining the default value of 50.

The hot and cold spot analysis was conducted using the Hot Spot Analysis (Getis – Ord Gi*) tool (Getis and Ord, 1992). The Getis-Ord Gi* statistic was chosen over more global statistics (e.g. The Global Moran’s I Index) to identify more local spatial clusters by finding local maxima. It was possible to compare of each feature with the surrounding neighbours (Getis & Ord, 1992). This analysis computes by a default a z-score (a measure of standard deviation) and p-value. A high z-score and small value of p indicate clustering of high values (hot spots), whilst low z-score with small p-value inform about spatial clustering of lower values (cold spots). Point data was exploited to detect the hot spot areas with significantly high salp densities and cold spots with their lower abundance. In the analysis we used inverse distance band in order to assure use of every point in the analysis in the decadal subsets.

The future boundary of temperature optimum for the distribution range of salps was established based on HadISST1 data from the Met Office Hadley Centre’s (Rayner et al., 2003). Mean distribution of surface temperatures during local summer (from December to the beginning of March) was created by mosaicking SST rasters for the years 1970 to 2001. The first boundary of salp occurence based on thermal preferences was determined by extracting the periphery of 0°C isotherm. Temperature range for salp occurrence was based on available literature data (Foxton, 1966; Ono & Moteki, 2016). The predicted boundary after a temperature increase of 1°C was extracted analogically from the periphery of 1°C isotherm. According to the previous studies, water temperature along the Western Antarctic Peninsula has increased by more than 1°C over the last 50 years (Whitehouse et al., 2008). We adopted a conservative approach and assumed that water temperature will increase by the next degree over the 50 years ahead. The maps presenting spatial distribution and abundances of *Salpa thompsoni* were created using ESRI^®^ ArcMap™ 10.5.1 software.

## Results

The extended collection resulted in 1872 samples, of which *Salpa thompsoni* was present in 1278. Our results from the long-term data series revealed significant annual variability in salp abundance. Through the late 1970s and early 1980s mean abundances of *S. thompsoni* were relatively low in researched areas, with the exceptions of the cruises in 1975/76 and 1983/84. After season 1987/88, with very low salps densities (0.9 ind/1000m^3^ ±1.26), the extremely high peak abundance occurred in the summer 1989/90, with the mean density 1408 ind/1000m^3^ (±2778.31). Although season 1994/95 was characterized by low salp abundances, their numbers began to increase over the following years (Fig. 2).

**Fig. 2.**
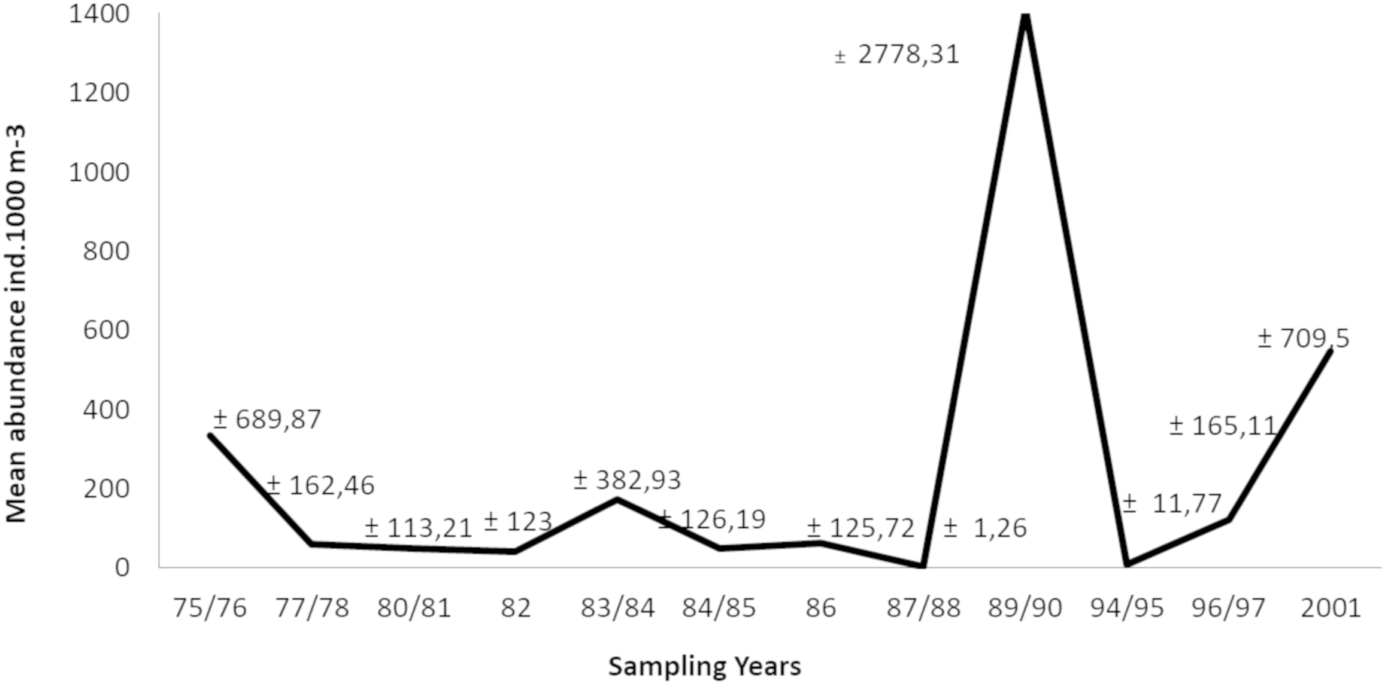
Mean abundance of *Salpa thompsoni* sampled each year (standard deviation values are given above the line).

Analysis based on in situ data also confirmed that the high abundance of salps was influenced by the sea ice concentration in the time of sampling (Kruskal-Wallis test H=33.98, p<0.0001). Sea ice played a crucial role in the Antarctic salp distribution, however significant differences were observed only between stations without ice cover (0) and with the presence of ice (1-3). There was no statistically significant difference between stations with a different degree of ice cover, which could affect presence of salps. Further analyzes have implied that *S. thompsoni* may occur in less favorable ice conditions (ice-cover around 1-3 degree) in smaller numbers. Interestingly, the probit model for the Sea Ice Index data, confirmed that the ice concentration during winter months (July and August) had a significant impact on the presence of salps in the following summer season (N=1311, χ2 =211.36, p<0.0001). The nonparametric Spearman’s rank-order revealed only weak correlations between selected environmental variables (bottom depth vs T rs=0.2, bottom depth vs sea ice rs=−0.25, and sea ice vs T rs=−0.16), therefore Generalized Additive Models (GAMs, Fig. 3) were tested independently and separately for above environmental variables. GAM fitted for sea surface temperature (F=5.3, p=0.0054) showed a uni-modal response, with the highest number of *S. thompsoni* registered in the water layers with temperatures between 1°C and −1°C (Fig. 3). GAM against salinity (F=5.8, p=0.0039) and bottom depth (F= 15.0, p<0.00001) did not give such a clear response. The highest numbers of salps seemed to be observed within waters with a salinity around 33.5-34.25. The same model used for the bottom depth showed the presence of the major peak, indicating preference for shallow shelf waters with bottom depth around 0-1000 m (Fig. 3).

**Fig. 3.**
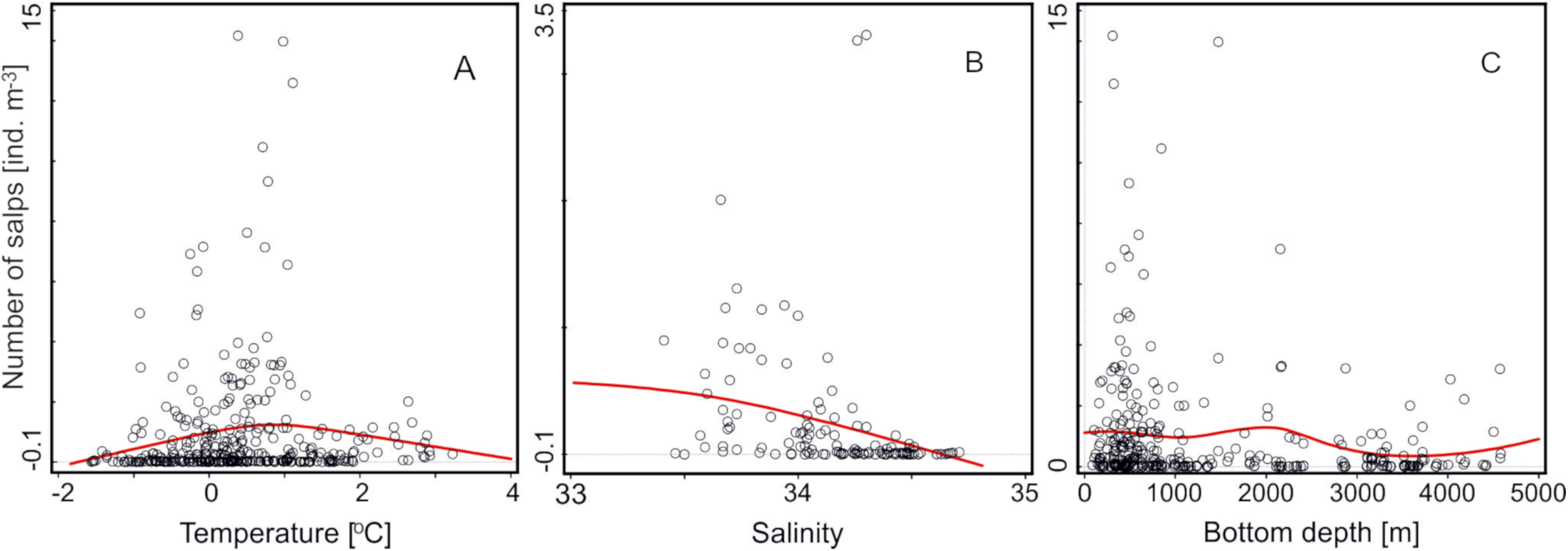
Generalized Additive Models (GAM) fitted for the number of salps against: (A) sea surface temperature (N=461), (B) salinity (N=212), and (C) bottom depth (N=1278).

### Hot spot Analysis

Despite the fact that sampling sites in the 1970’s were spread along the West Antarctic Peninsula region, near the South Shetland Islands, around South Orkney Islands in the Atlantic sector of the Southern Ocean, statistically significant hot spots were located mostly near Elephant Island, where ice concentration was degree 0 (Z-score > 1.67 and p value < 0.05). In the 1980’s a hot spot was registered further to the south. Regardless of the similarity between sampling coverages of WAP area, hot spots (Z-score > 1.7 and p value < 0.05) were placed in regions of the Biscoe Islands, as well as Anvers Island and through the Bransfield Strait (Fig. 4). Significant high densities of salps, assuming 99% confidence, were still found in the vicinity of Elephant Island. The stations with high abundance of salps were in the areas with ice coverage classified as 1. Sampling stations in seasons 1994/95, 1996/97 and 2001 were clustered around the Elephant Island region. This region showed *S. thompsoni* hot spots, with a high Z-score (up to 7.89) and p value <0.05.

**Fig 4.**
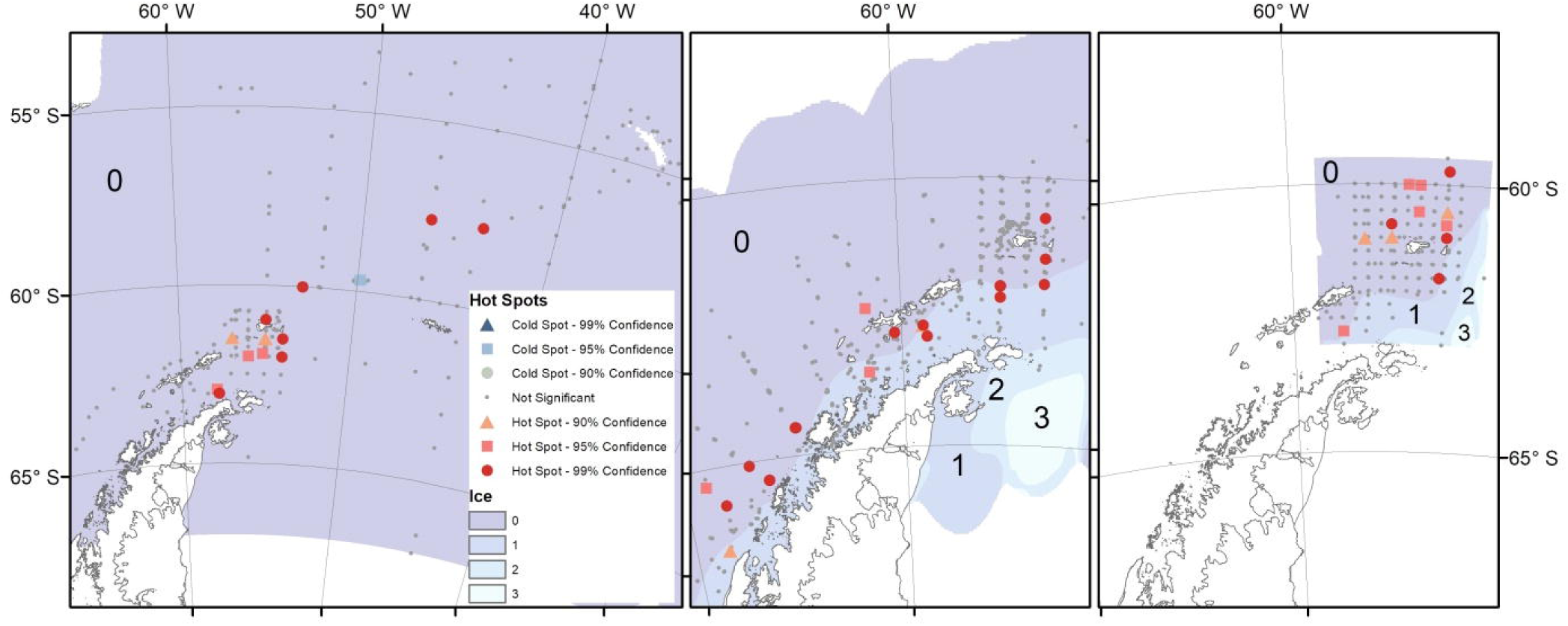
Hot spots of *Salpa thompsoni* distribution in decadal time series: (A) ‘70s, (B) ‘80s and (C) ‘90s (in the background - the interpolated ice concentration for the respective time series. Cold spotsmarked: in grey circles - 90% confidence, blue squares - 95% confidence, dark blue triangles - 99 % confidence; hot spots marked: in orange triangles - 90% confidence, dark orange sqares - 95% confidence, red circles – 99% confidence; different sea ice degrees marked in: violet – without sea ice, dark blue-1, sky blue-2, light blue-3).

## Discussion

Our results revealed significant annual variability of *Salpa thompsoni* due to differences in bottom depth and environmental conditions e.g. water temperature or ice condition prevailing during and before sampling years. Presented data also indicated that the crucial months for salps were July and August when the lack of sea ice presumably allows these animals to form more vast and condensed aggregations during the following summer periods. Also, the previous studies revealed that salps were most frequently present in the ice-free regions (Atkinson, Siegel, Pakhomov & Rothery, 2004; Chiba, Ishimaru, Hosie & Wrigh, 1999; Lee, Pakhomov, Atkinson & Siegel, 2010; Mackey et al., 2011; Ross et al., 2014; Steinberg et al., 2015), which our results initially confirmed. Although, our further analysis showed that *S. thompsoni* may be present even with less suitable ice conditions, 1-2 ice degree, which means that they might be able to exist in colder environment. Used GAM model, revealed untypical salp preferences, different than previously noted, showing that the highest salp number was located in the shallow shelf waters (bottom depths 0-1000 m), with a low temperature ([-1] – 1°C), and salinity of 33.5 – 34.25 (Fig. 3). Our observation was contrary to the conclusions of Atkinson, Siegel, Pakhomov & Rothery (2004) who believed that cold shelf water is a place occupied by krill but will not be preferred by Antarctic salps. This finding may suggest possible shifts of salp densities closer to the cold continent, or could be partially explained by the dynamics of the ACC water masses. The ACC pumps warm (>1.5°C), salty (34.65–34.7), and nutrient-rich Circumpolar Deep Water (CDW) onto the continental shelf below 200 m (Klinck, Hofmann, Beardsley, Salihoglu & Howard, 2004), and may moves salps towards the Antarctic continent and the surrounding islands to the Bransfield Strait areas (Tokarczyk, 1987), where they can be found in high abundances.

The Hot Spots Analysis in three decadal time periods - ‘70s, ‘80s, and ‘90s - were confirmed that the *Salpa thompsoni* distribution has been located mostly around Elephant Island. Our dataset from ‘70s and ‘90s showed that their greatest numbers were observed mostly within open ocean as well as around the AP area, however, in the ‘80s *S. thompsoni* hot spots were located evenly within Islands around Bransfield and Gerlache Straits, as well as far south near the Bellingshausen Sea, where sea ice was present and water temperature was lower. The greatest salp number was recorded between 65-70°S in the presence of ice-cover and water temperature below 0°C, which can also confirm previous observations by Ono & Moteki (2013), and may suggest that *S. thompsoni* is able to adapt to less suitable conditions in colder, shelf waters. S. *thompsoni* generally occurs in warm water masses > 0 °C, between 45-55°S (Foxton, 1966). Results of this study showed that a high number of mature individuals could be observed even south of the Southern Boundary (SB). Our results, together with the study presented by Ono & Moteki (2013), has revealed that *S. thompsoni* appeared across the SB of the ACC, and suggests that presumably it is not a barrier preserving Antarctic waters from *S. thompsoni* southward migration. A number of genetic studies further proved that salps are highly flexible (Batta-Lona, Maas, O’Neill, Wiebe & Bucklin, 2017; Goodall-Copestake, 2016; Jue et al., 2016), capable of adaptation to the changing environmental conditions within different parts of the Southern Ocean. The comprehensive data about modification of salps/krill population were presented by Ross et al. (2014). They showed that krill abundance had decreased on the northern lines and had shifted 200 km to the south. If the trend of shrinking sea ice continues, there is a certain risk that within two decades, krill may completely disappear from the northern region of the Southern Ocean. Herein presented result expands foregoing knowledge about the dynamics of Antarctic salp populations and demonstrates missing predictions of their future distribution. In our study, we have presented inference about possible future salps distribution modification with the assumption that the water temperature will increase by 1°C, and results showed that the *S. thompsoni* range could move southwards by an average of 200 km.

Taking into account climate fluctuations and temperature trends, we tracked the movement of the temperature boundary for salps for a hypothetical situation of ocean warming by 1°C for the next 50 years. The comparison of the locations of isotherms 0°C and 1°C revealed that massive blooms of *S. thompsoni* possible may shift their distribution southward e.g. into the Weddell Sea and closer to the Antarctic continent. Such temperature rise would enlarge their habitat area by nearly 530 000 km^2^, what possibly may exclude Antarctic krill and consequently lead to changes in the functioning of the Antarctic ecosystem (Fig. 5).

**Fig. 5.**
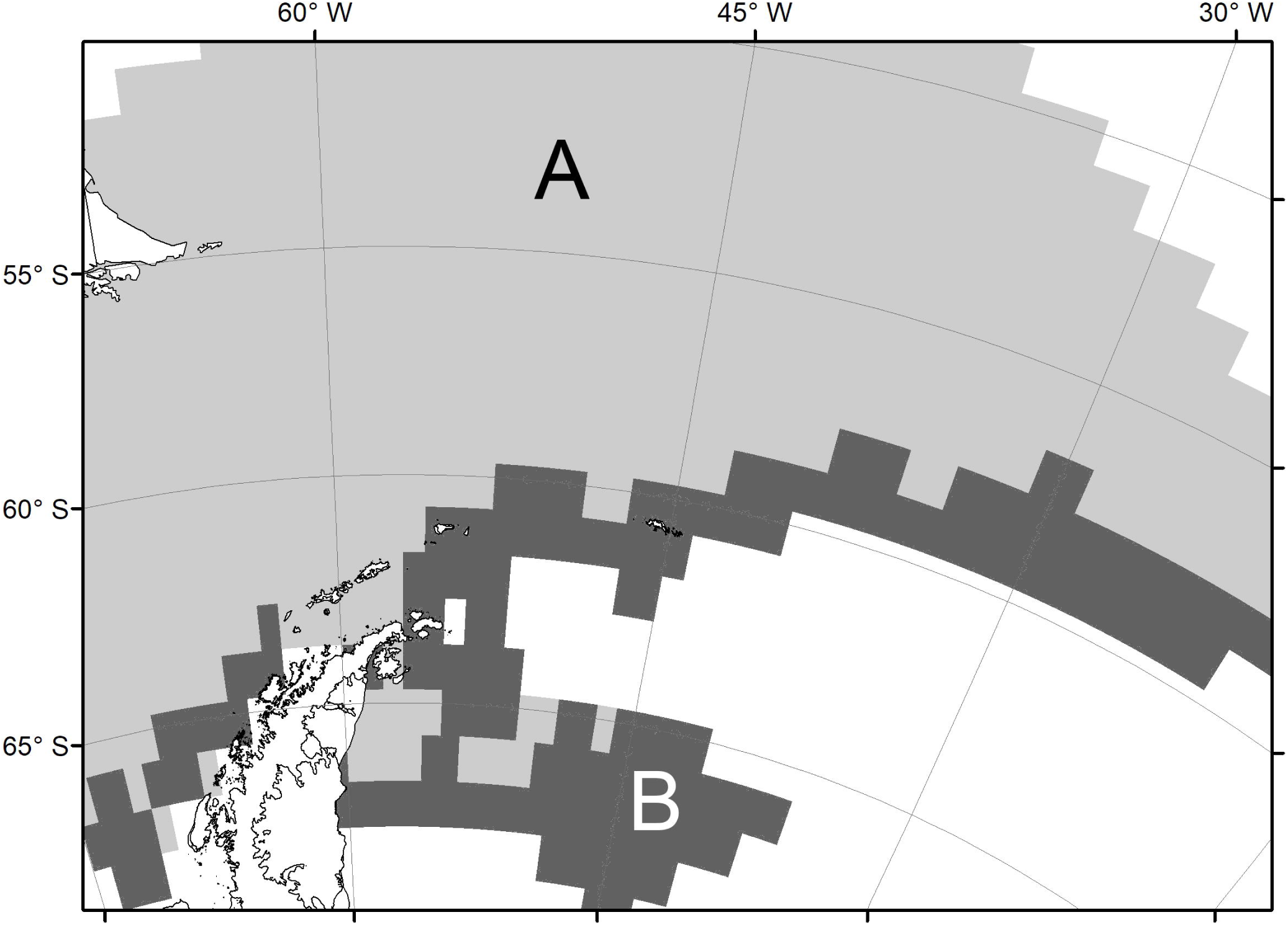
Boundary of *Salpa thompsoni* temperature optimum: in the studied year (A - marked in light gray) and prognosis on its change after 1°C temperature rise (B - marked in dark gray).

## Acknowledgments

We want to express our gratitude to the crews and scientists who have collected and analysed these samples over the last 3 decades, and provided this data in a useable format. We are grateful to Prof. Evgeny Pakhomov from University of British Columbia, M.Sc. Natasha Waller from Australian Antarctic Division for their willingness and help in improving this manuscript. We would like to thank Dr. Adrian Zwolicki from the University of Gdansk for his advice concerning statistical analyses. The authors declare that they have no conflict of interest.

